# Selection signatures in worldwide sheep populations

**DOI:** 10.1101/001453

**Authors:** Maria-Ines Fariello, Bertrand Servin, Gwenola Tosser-Klopp, Rachel Rupp, Carole Moreno, International Sheep Genomics Consortium, Magali SanCristobal, Simon Boitard

**Affiliations:** Génétique, Physiologie et Systémes d’Élevage, INRA & ENVT & ENSAT, Castanet-Tolosan, France; IMERL, Facultad de Ingeniera, Universidad de la Repblica, Montevideo, Uruguay; Bioinformatics Unit, Institut Pasteur, Montevideo, Uruguay; www.sheephapmap.org; Génétique Animale et Biologie Intégrative, INRA & AgroParisTech, Jouy-en-Josas, France; Origine, Structure et Evolution de la Biodiversité, Museum National d’Histoire Naturelle & EPHE & CNRS, Paris, France

## Abstract

The diversity of populations in domestic species offers great opportunities to study genome response to selection. The recently published Sheep HapMap dataset is a great example of characterization of the world wide genetic diversity in sheep. In this study, we re-analyzed the Sheep HapMap dataset to identify selection signatures in worldwide sheep populations. Compared to previous analyses, we made use of statistical methods that (i) take account of the hierarchical structure of sheep populations, (ii) make use of linkage disequilibrium information and (iii) focus specifically on either recent or older selection signatures. We show that this allows pinpointing several new selection signatures in the sheep genome and distinguishing those related to modern breeding objectives and to earlier post-domestication constraints. The newly identified regions, together with the ones previously identified, reveal the extensive genome response to selection on morphology, color and adaptation to new environments.

## Introduction

Domestication of animals and plants has played a major role in human history. With the advance of high-throughput genotyping and sequencing technologies, the analysis of large datasets in domesticated species offers great opportunities to study genome evolution in response to phenotypic selection [1]. The sheep was one of the first grazing animals to be domesticated [2] in part due to its manageable size and an ability to adapt to different climates and diets with poor nutrition. A large variety of breeds with distinct morphology, coat color or specialized production (meat, milk or wool) were subsequently shaped by artificial selection. Since the release of the 50 K SNP array [3], it is now possible to scan genetic diversity in sheep in order to detect loci that have been involved in these various adaptive selection events. The Sheep HapMap dataset, which includes 50 K genotypes for 3000 animals from 74 breeds with diverse world-wide origins, provides a considerable resource for deciphering the genetic bases of phenotype diversification in sheep. In the first analysis of this dataset [4], the authors looked for selection by computing a global *F_ST_* among the 74 breeds at all SNP in the genome. They identified 31 genome regions with extreme differentiation between breeds, which included candidate genes related to coat pigmentation, skeletal morphology, body size, growth, and reproduction. Further studies took advantage of the Sheep HapMap resource to detect genetic variants associated with pigmentation [5], fat deposition [6], or microphtalmia disease [7]. An other study [8] performed a genome scan for selection focused on American synthetic breeds, using an *F_ST_* approach similar to that in [4].

The 74 breeds of the Sheep HapMap dataset have a strong hierarchical structure, with at least 3 distinct differentiation levels: an inter-continental level (e.g. European breeds vs Asian breeds), an intracontinental level (e.g. Texel vs Suffolk European breeds), and an intra-breed level (e.g. German Texel vs Scottish Texel flocks). Recent studies [9–12] showed that, when applied to hierarchically structured data sets, *F_ST_* based genome scans for selection may lead to a large proportion of false positives (neutral loci wrongly detected as under selection) and false negatives (undetected loci under selection). Besides, the heterogeneity of effective population size among breeds implies that some breeds are more prone to contribute large locus-specific *F_ST_* values than others [10]. Apart from these statistical considerations, merging populations with various degrees of shared ancestry can limit our understanding of the selective process at detected loci. Indeed, the regions pointed out in [4] can be related to either ancient selection, as the poll locus which has likely been selected for thousands of years, or fairly recent selection, as the myostatin locus which has been specifically selected in the Texel breed. But in most situations the time scale of adaptation cannot be easily determined.

Another limit of genome scans for selection based on single SNP *F_ST_* computations is that they do not sufficiently account for the very rich linkage disequilibrium information, even when the single SNP statistics are combined into windowed statistics. Recently, we proposed a new strategy to evaluate the haplotype differentiation between populations [13]. We showed that using this approach greatly increases the detection power of selective sweeps from SNP chip data, and also enables to detect soft or incomplete sweeps. These latter selection scenarios are particularly relevant in breeding populations, where selection objectives have likely varied along time and where the traits under selection are often polygenic.

In this study we provide a new genome scan for selection based on the Sheep HapMap dataset, where we distinguish selective sweeps within and between 7 broad geographical groups. The within group analysis aims at detecting recent selection events related to the diversification of modern breeds. It is based on the single marker FLK test [10] and on its haplotypic extension hapFLK [13]. The FLK test is an extension of the Lewontin and Krakauer (LK) test [14] that accounts for population size heterogeneity and for the hierarchical structure between populations. As the LK test, the FLK test computes a global *F_ST_* for each SNP, but allele frequencies are first rescaled using a population kinship matrix *F* . This matrix, which is estimated from the observed genome wide data, measures the amount of genetic drift that can be expected, under neutral evolution, along all branches of the population tree. With this rescaling, allele frequency differences are typically down-weighted if they are obtained with small populations, or populations that diverged a long time ago. The between group analysis focuses on older selection events and is only based on FLK. Overall, we confirmed 19 of the 31 sweeps discovered in [4], while providing more details about the past selection process at these loci. We also identified 71 new selection signatures, with candidate genes related to coloration, morphology or production traits.

## Results and discussion

We detected selection signatures using methods that aim at identifying regions of outstanding genetic differentiation between populations, based either on single SNP, FLK [10], or haplotype, hapFLK [13], information. These methods have optimal power when working on closely related populations so we separately analyzed seven groups of breeds, previously identified as sharing recent common ancestry [4] and corresponding to geographical origins of breeds. Before performing genome scans for selection signatures, we studied the population structure of each group to identify outlier animals as well as admixed and strongly bottlenecked populations, using both PCA and model-based approaches [15,16]. hapFLK was found to be robust to bottlenecks or moderate levels of admixture, but these phenomena may affect the detection power so we preferred to minimize their influence by removing suspect animals or populations. Details of these corrections are provided in the methods section. The final composition of population groups are given in Table 1.

**Table 1.**
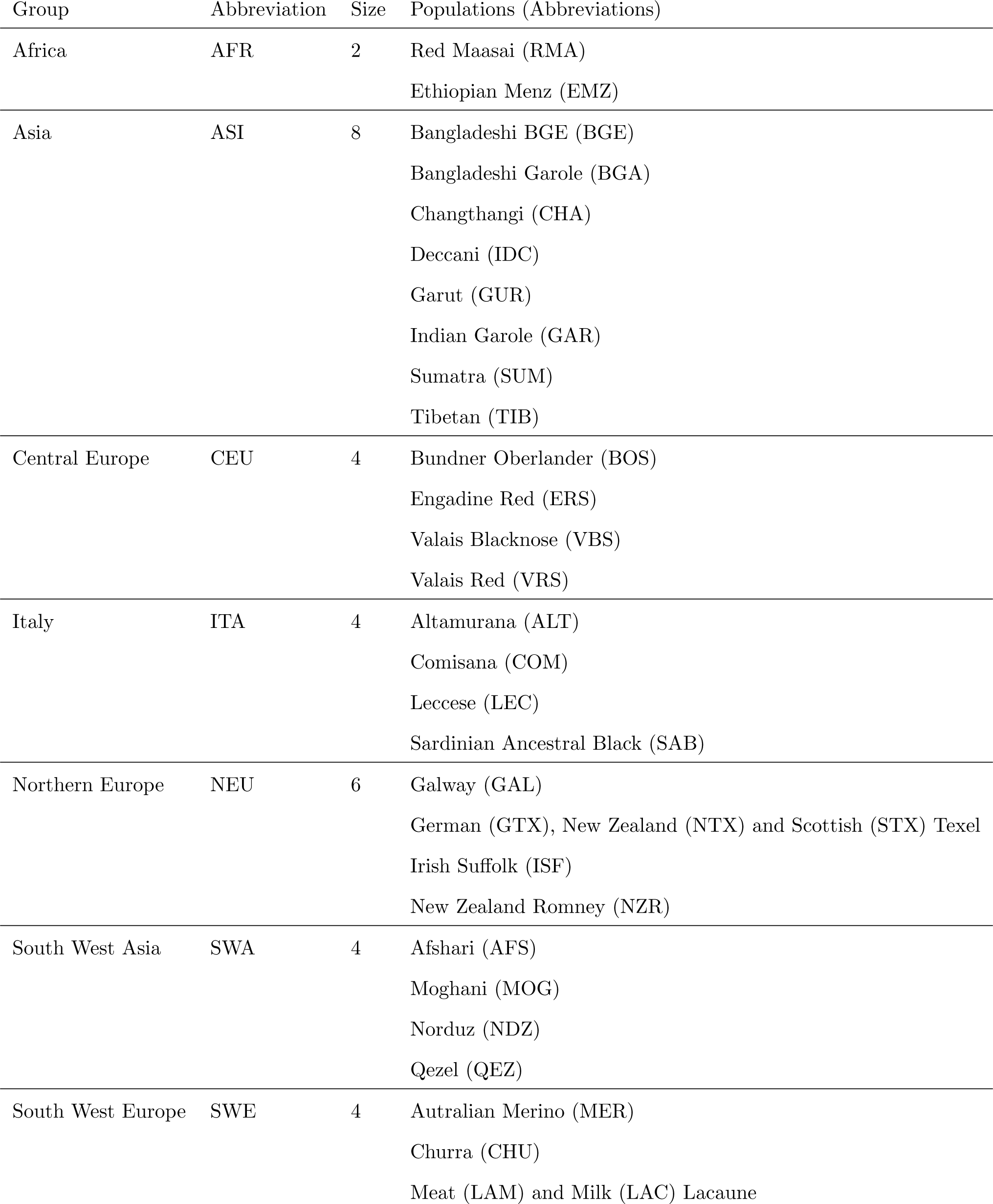
Population groups from the Sheep HapMap dataset used for the detection of selection signatures

| Group | Abbreviation | Size | Populations (Abbreviations) |
| --- | --- | --- | --- |
| Africa | AFR | 2 | Red Maasai (RMA)<br>Ethiopian Menz (EMZ) |
| Asia | ASI | 8 | Bangladeshi BGE (BGE)<br>Bangladeshi Garole (BGA)<br>Changthangi (CHA)<br>Deccani (IDC)<br>Garut (GUR)<br>Indian Garole (GAR)<br>Sumatra (SUM)<br>Tibetan (TIB) |
| Central Europe | CEU | 4 | Bundner Oberlander (BOS)<br>Engadine Red (ERS)<br>Valais Blacknose (VBS)<br>Valais Red (VRS) |
| Italy | ITA | 4 | Altamurana (ALT)<br>Comisana (COM)<br>Leccese (LEC)<br>Sardinian Ancestral Black (SAB) |
| Northern Europe | NEU | 6 | Galway (GAL)<br>German (GTX), New Zealand (NTX) and Scottish (STX) Texel<br>Irish Suffolk (ISF)<br>New Zealand Romney (NZR) |
| South West Asia | SWA | 4 | Afshari (AFS)<br>Moghani (MOG)<br>Norduz (NDZ)<br>Qezel (QEZ) |
| South West Europe | SWE | 4 | Australian Merino (MER)<br>Churra (CHU)<br>Meat (LAM) and Milk (LAC) Lacaune |

### Overview of selected regions

An overview of selection signatures on the genome across the different groups is plotted in Figure 1 and a detailed description is provided in Table 2. Detected regions were typically a few megabases long and included from 1 to 196 genes, with a median of 15 genes. However, in many regions strong functional candidate genes were found very close to the position with lowest p-value, typically among the two closest genes from this position. These genes are reported in Table 2, as well as a few other functional candidates with less statistical evidence but strong prior knowledge from the literature. We found 41 selection signatures with hapFLK and 26 with FLK, although we allowed a slightly higher false discovery rate for FLK than hapFLK (10% vs 5%). This result was consistent with a higher power for hapFLK than FLK, as already shown in [13].

**Figure 1.**
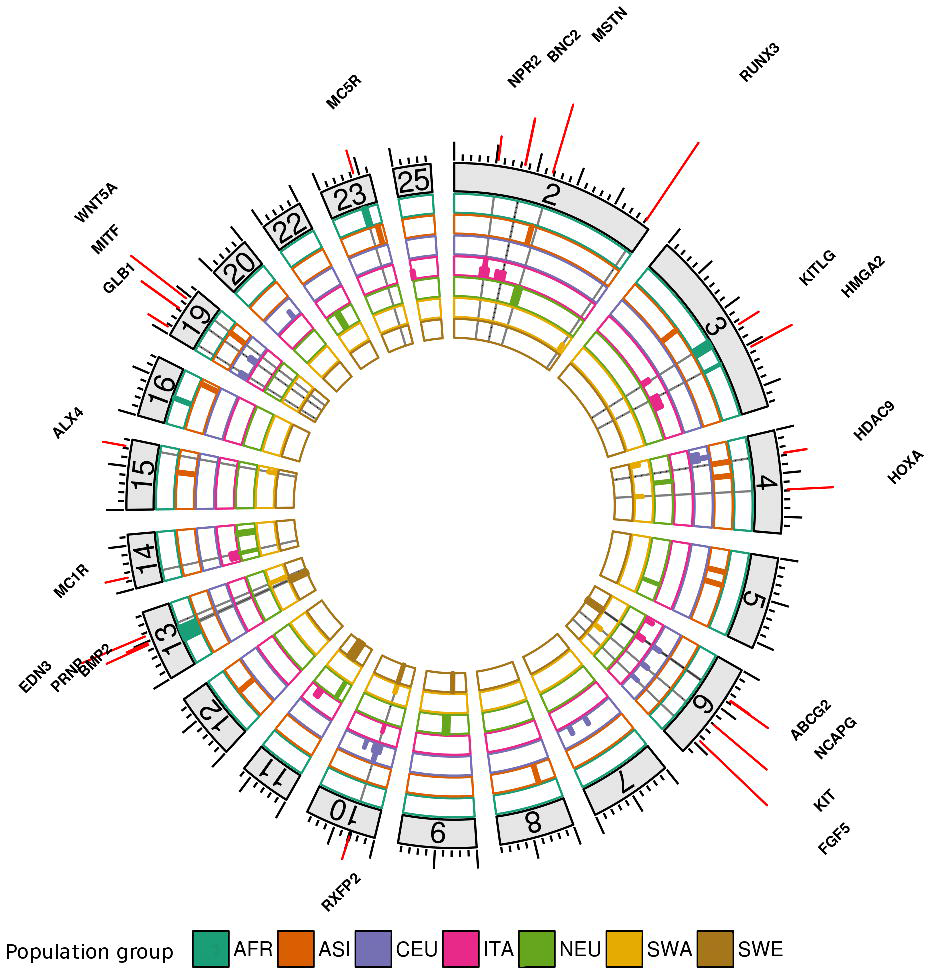
Localization of selection signatures identified in 7 groups of populations. Candidate genes are indicated above their genomic localization. Only chromosomes harboring selection signatures are plotted.

**Table 2.**
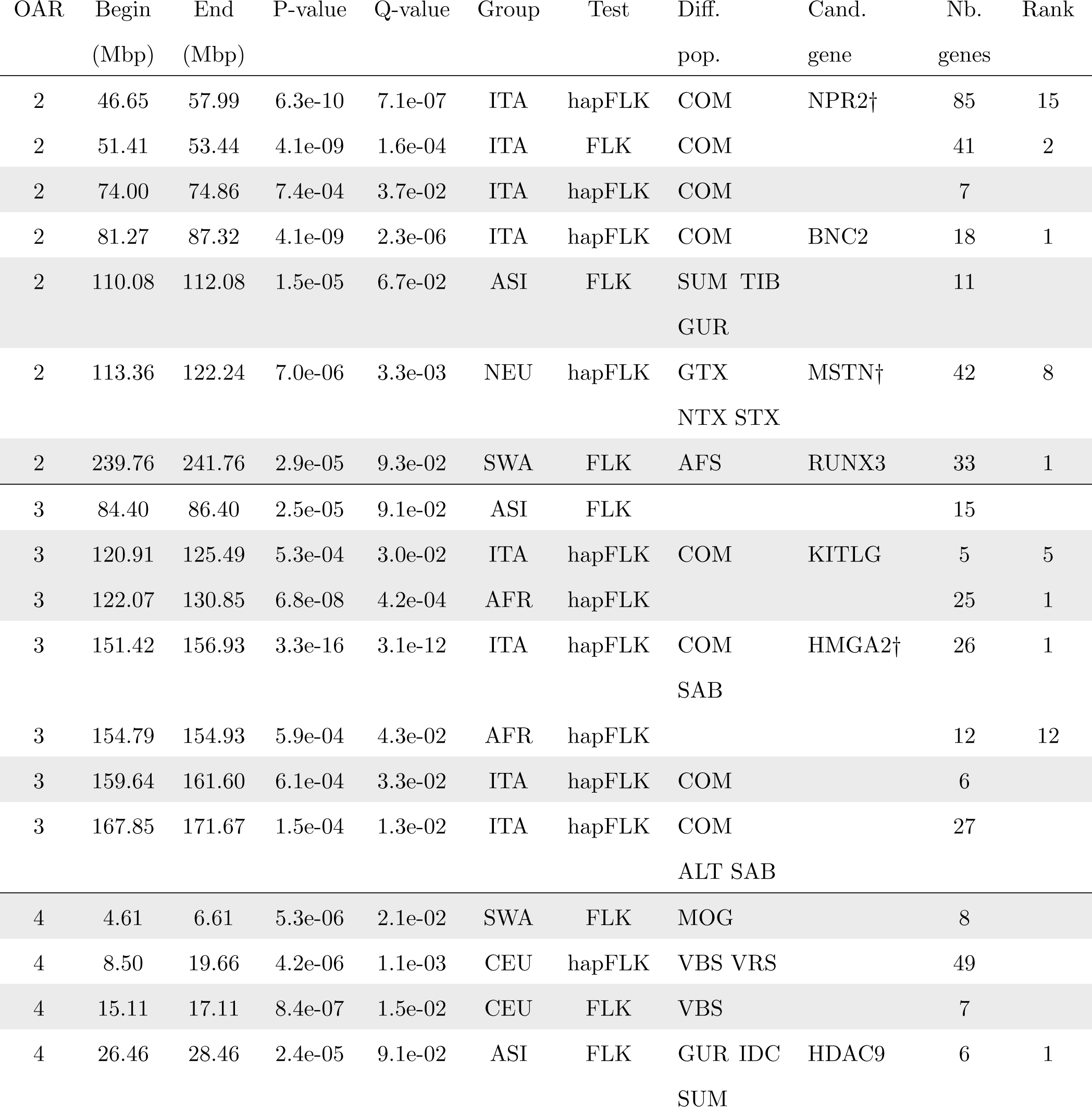
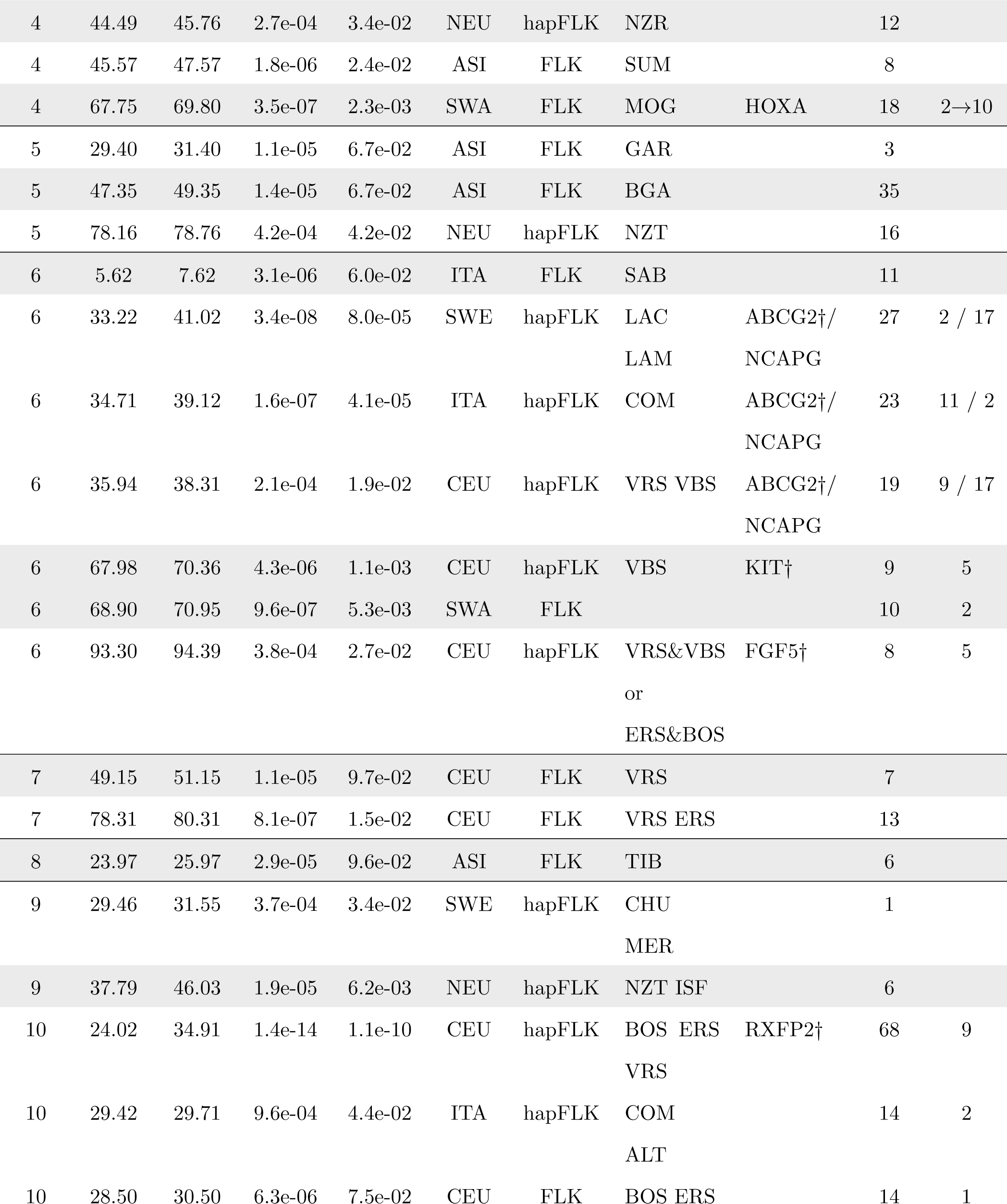
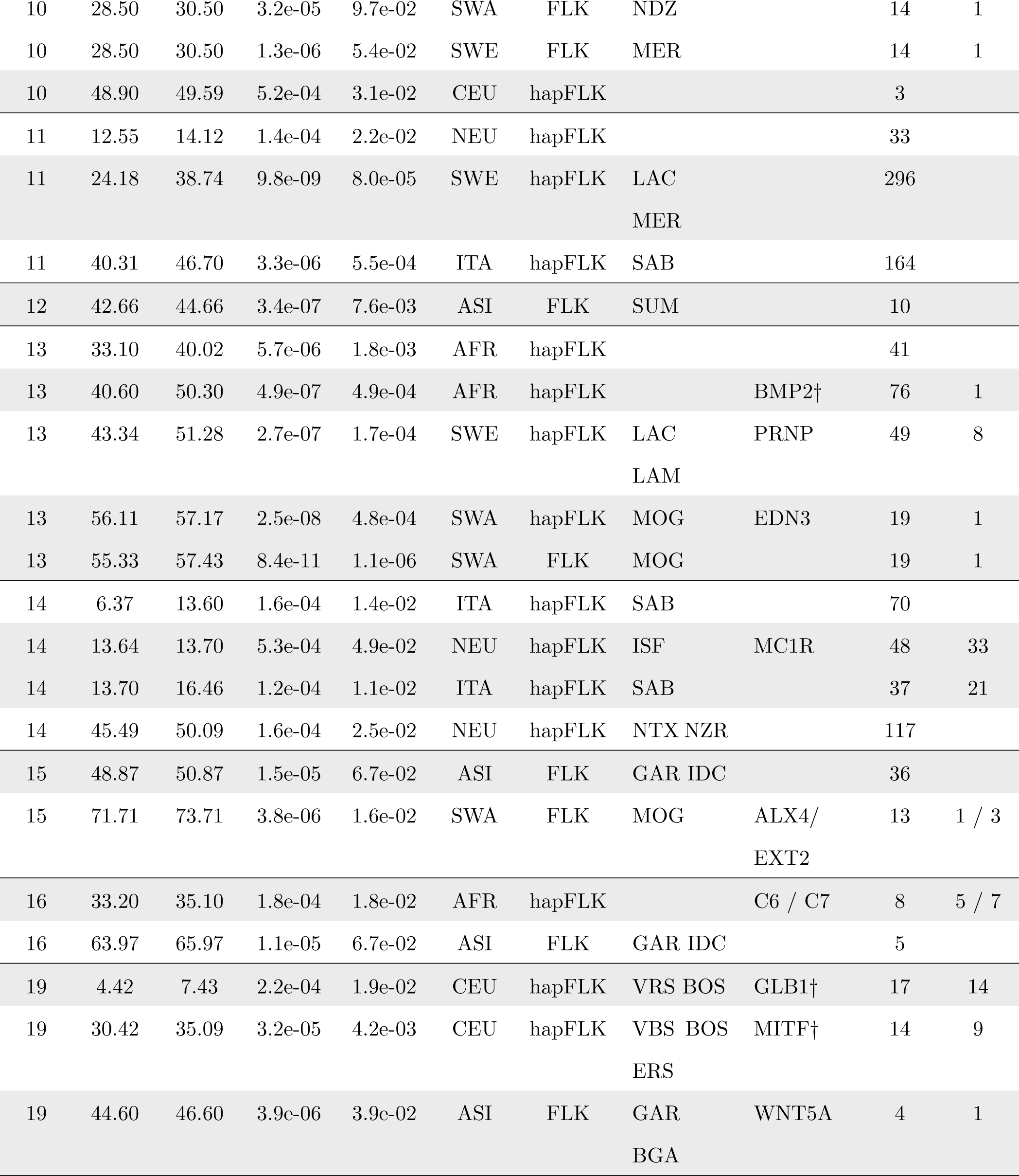
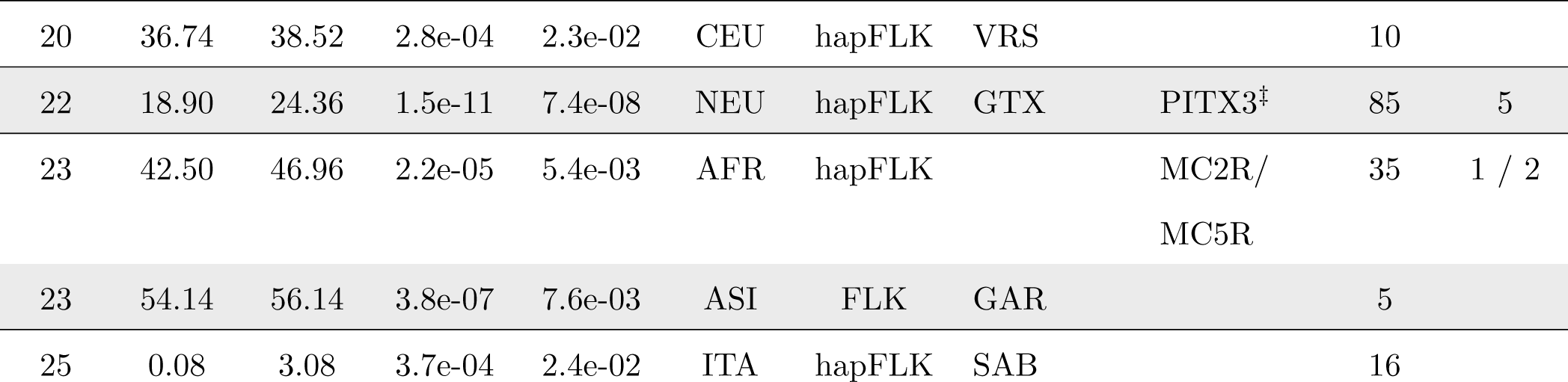
Selection signatures in the 7 geographical groups. Regions identified with the hapFLK or FLK test, with the corresponding population group and most differentiated populations (except for the AFR group). Full names of groups and populations are given in Table 1. The number of genes included in each region and the rank of candidate genes within the region is also provided. Overlapping regions in different groups or with different tests are grouped by background color.†: signatures of selection previously identified [4].‡: this outlying region is not due to evolutionary processes (see details in the main text).

Four regions were found with both the single SNP and the haplotype test and harbor strong candidate genes: NPR2, KIT, RXFP2 and EDN3 (Table 2). The overlap was thus small, illustrating that the two tests tend to capture different signals. In particular, hapFLK will fail to detect ancient selective sweeps, for which the mutation-carrying haplotype is small and not associated with many SNP on the chip. On the contrary, single SNP tests will fail to capture selective sweeps when a single SNP is not in high LD with the causal mutation. They will also fail if the selected mutation is only at intermediate frequency but is associated to a long haplotype, in contrast with hapFLK.

Six regions were detected in more than one group of breeds. They all contained strong candidate genes (Table 2). Three of these genes are related to coat color (KIT, KITLG and MC1R), and could correspond to independent selection events (see discussion below). One region harbors a gene (RXFP2) for which polymorphisms have been shown to affect horn size and polledness in the Soay [17] and Australian Merino [18]. We detected this region in 4 different groups and in all of them the highest FLK value was found to be very close to RXFP2 (Figure S8). This provides clear evidence that selection in this region is related to RXFP2, consistent with previous selection signatures detected by comparing specifically horned and polled breeds (Figure 6 in [4]). However, we note that the signatures of selection in this region exhibit different patterns among groups. The signal is very narrow in the SWE and SWA groups, and is in fact not detected by the hapFLK test, whereas it affects a large genome region in the CEU group where it is detected by hapFLK. In the ITA group, the FLK statistics do not reach significance, and the hapFLK signal is not high (minimum q-value of 0.04). Overall, the selection signatures suggest that selection on RXFP2, most likely due to selection on horn phenotypes, was carried out worldwide at different times and intensities. Another region harbors the HMGA2 gene, involved in selection for stature in dogs [19]. The last region includes two interesting candidate genes : ABCG2, which has been associated to a strong QTL for milk production in cattle [20], and NCAPG, which has been associated to fetal growth [21] and calving ease [22] in cattle and which is located in several selection signatures in this species [23–26]. In our analysis, populations with a selection signature in this region belong to three European groups (SWE, ITA and CEU) and our results suggest that selection in these different groups might imply distinct genes (Table 2).

In the paper presenting the Sheep HapMap dataset [4], 31 selection signatures were found, corresponding to the 0.1% highest single SNP *F_ST_* . Using FLK and hapFLK, we confirmed signatures of selection for 10 of these regions. Considering the two analyses were performed on the same dataset, this overlap can be considered as rather small. Two reasons can explain this.

First, the previous analysis was based on the *F_ST_* statistic. Although this statistic is commonly used for selection scans, it is prone to produce false positives when the population tree harbors unequal branch lengths (*i.e.* unequal effective population sizes) [10]. In particular, strongly bottlenecked breeds will contribute high *F_ST_* values preferentially even under neutral evolution, because their smaller effective population size implies a larger variance of allele frequencies. With *FLK* and *hapFLK*, *F_ST_* values between populations are rescaled using branch lengths, so populations with long branch lengths will not contribute more than others [13]. In fact they will tend to contribute less, as the statistical power to distinguish selective effects from drift effects is naturally lower in populations where drift is larger.

Second, the previous analysis was performed using all breeds at the same time. It is therefore possible that some of these regions correspond to differentiation between groups of breeds rather than within groups. To investigate this question, we performed a genome scan for selection between the ancestors of the seven population groups using the FLK statistic computed on their estimated allele frequencies [10]. We did not include SNP lying in regions detected within groups since selection biases their estimated ancestral allele frequencies. The population tree was reconstructed using SNP for which we have unambiguous ancestral allele information (Figure S9). The tree is decomposed into two main lineages, one for European breeds and one for Asian and African breeds. The African group exhibits a slightly higher branch length. We note, however, that this could be due to ascertainment bias of SNP on the SNP array.

This led to the identification of 23 new selection signatures (Figure 2 and Table 3), 9 of them being common to the analysis of [4]. Overall, combining the scans for recent and ancestral selection, we failed to replicate 12 of the regions in [4].

**Figure 2.**
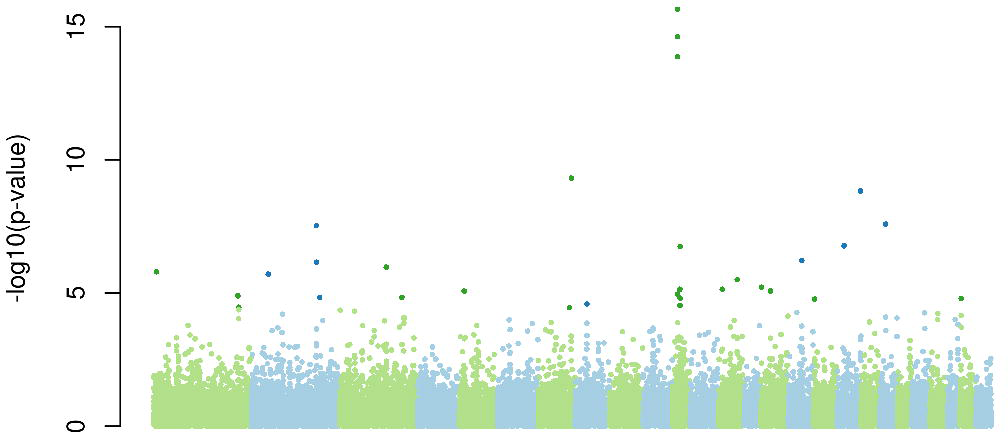
Genome scan for selection signature in ancestral populations of the geographical groups. Significant SNP at the 5% FDR level are plotted in darker color.

**Table 3.** Selection signatures in ancestral populations. SNP with significant FLK value at the 5% FDR level, with estimated allele frequencies in all ancestral groups. The number of genes included in each region (1Mb up-or-downstream the position) and the rank of candidate genes within the region is also provided. †: signatures of selection previously identified [4].

| OAR | position | Estimated ancestral allele frequencies |  |  |  |  |  |  | P-value | Q-value | Cand. gene | Nb. genes | Rank |
| --- | --- | --- | --- | --- | --- | --- | --- | --- | --- | --- | --- | --- | --- |
|  |  | AFR | ASI | SWA | NEU | CEU | ITA | SWE |  |  |  |  |  |
| 1 | 7192190 | 0.15 | 0.08 | 0.16 | 0.55 | 0.69 | 0.04 | 0.38 | 1.7e-06 | 5.3e-03 | TRPM8 | 19 | 8 |
| 1 | 237070498 | 0.87 | 0.95 | 0.91 | 0.48 | 0.24 | 0.77 | 0.35 | 1.4e-05 | 2.5e-02 | GYG1 | 16 | 5 |
| 1 | 239424807 | 0.46 | 0.68 | 0.06 | 0.21 | 0.15 | 0.11 | 0.17 | 3.4e-05 | 4.8e-02 |  | 9 |  |
| 1 | 239491620 | 0.53 | 0.41 | 0.94 | 0.86 | 0.93 | 0.93 | 0.88 | 4.3e-05 | 5.6e-02 |  | 9 |  |
| 2 | 45500785 | 0.43 | 0.91 | 0.23 | 0.76 | 0.87 | 0.87 | 0.93 | 2.2e-06 | 6.4e-03 | LPL | 6 | 3 |
| 2 | 182607165 | 0.99 | 0.97 | 0.18 | 0.64 | 0.73 | 0.83 | 0.64 | 3.4e-08 | 1.8e-04 | INSIG2 | 10 | 3 |
| 2 | 182672296 | 0.99 | 0.94 | 0.32 | 0.90 | 0.86 | 0.89 | 0.81 | 7.7e-07 | 2.8e-03 |  | 10 |  |
| 2 | 192231314 | 0.59 | 0.93 | 0.36 | 0.96 | 0.89 | 0.81 | 0.95 | 1.6e-05 | 2.8e-02 |  | 8 |  |
| 3 | 132478420 | 0.24 | 0.89 | 0.18 | 0.93 | 0.81 | 0.84 | 0.82 | 1.2e-06 | 3.9e-03 | HOXC † | 54 | 1→9 |
| 3 | 180860403 | 0.71 | 0.53 | 0.28 | 0.82 | 0.31 | 0.12 | 0.13 | 1.7e-05 | 2.8e-02 |  | 22 |  |
| 5 | 15522700 | 0.68 | 0.63 | 0.92 | 0.27 | 0.76 | 0.99 | 0.78 | 9.8e-06 | 2.0e-02 |  | 51 |  |
| 7 | 89519883 | 0.63 | 0.61 | 0.19 | 0.89 | 0.18 | 0.60 | 0.95 | 6.1e-10 | 5.2e-06 | TSHR † | 6 | 3 |
| 8 | 31748642 | 0.84 | 0.93 | 0.94 | 0.16 | 0.63 | 0.47 | 0.19 | 2.8e-05 | 4.1e-02 | PREP † | 6 | 1 |
| 11 | 18248852 | 0.35 | 0.32 | 0.82 | 0.64 | 0.94 | 0.96 | 0.92 | 1.3e-05 | 2.5e-02 | NF1 † | 23 | 1 |
| 11 | 18325488 | 0.87 | 0.93 | 0.00 | 0.35 | 0.04 | 0.03 | 0.04 | 3.3e-16 | 7.2e-12 |  | 24 | 4 |
| 11 | 18335747 | 0.87 | 0.93 | 0.00 | 0.35 | 0.04 | 0.03 | 0.04 | 3.3e-16 | 7.2e-12 |  | 22 | 4 |
| 11 | 18433474 | 0.87 | 0.93 | 0.02 | 0.35 | 0.07 | 0.02 | 0.05 | 3.8e-15 | 5.4e-11 |  | 22 | 1 |
| 11 | 18440783 | 0.78 | 0.93 | 0.02 | 0.34 | 0.07 | 0.02 | 0.05 | 2.0e-14 | 2.2e-10 |  | 22 | 1 |
| 11 | 25704651 | 0.97 | 0.96 | 0.97 | 0.42 | 0.94 | 0.94 | 0.96 | 8.5e-06 | 1.9e-02 |  | 73 |  |
| 11 | 26284826 | 0.99 | 0.97 | 0.94 | 0.38 | 0.93 | 0.95 | 0.79 | 3.2e-05 | 4.6e-02 |  | 100 |  |
| 11 | 26571629 | 0.92 | 0.94 | 0.98 | 0.29 | 0.89 | 0.88 | 0.86 | 1.8e-05 | 2.8e-02 |  | 115 |  |
| 11 | 26872280 | 0.78 | 0.71 | 0.93 | 0.15 | 0.89 | 0.90 | 0.90 | 2.2e-07 | 9.5e-04 |  | 111 |  |
| 13 | 12120674 | 0.29 | 0.84 | 0.97 | 0.91 | 0.97 | 0.92 | 0.84 | 7.7e-06 | 1.8e-02 | GATA3 | 6 | 1 |
| 13 | 62857560 | 0.52 | 0.62 | 0.65 | 0.98 | 0.67 | 0.92 | 0.36 | 3.6e-06 | 9.7e-03 | ASIP † | 32 | 12 |
| 15 | 3706790 | 0.71 | 0.22 | 0.96 | 0.28 | 0.27 | 0.34 | 0.21 | 6.8e-06 | 1.7e-02 |  | 4 |  |
| 15 | 29856310 | 0.98 | 0.99 | 0.99 | 0.47 | 0.92 | 0.95 | 0.96 | 9.8e-06 | 2.0e-02 |  | 35 |  |
| 16 | 38696505 | 0.95 | 0.98 | 0.95 | 0.99 | 0.68 | 0.31 | 0.30 | 6.8e-07 | 2.7e-03 | PRLR † | 18 | 2 |
| 17 | 4867509 | 0.91 | 0.95 | 0.85 | 0.54 | 0.18 | 0.58 | 0.17 | 1.8e-05 | 2.8e-02 | TMEM154 | 9 | 1 |
| 18 | 19342316 | 0.90 | 0.79 | 0.67 | 0.35 | 0.75 | 0.10 | 0.09 | 1.9e-07 | 9.3e-04 | ACAN † | 31 | 4 |
| 18 | 66470371 | 0.99 | 0.97 | 0.90 | 0.90 | 0.18 | 0.04 | 0.08 | 1.9e-09 | 1.3e-05 | TRAF3 | 28 | 5 |
| 20 | 17381047 | 0.24 | 0.61 | 0.97 | 0.98 | 0.93 | 0.99 | 0.91 | 3.1e-08 | 1.8e-04 | VEGFA † | 48 | 1 |
| 25 | 7517270 | 0.95 | 0.94 | 0.93 | 0.14 | 0.27 | 0.57 | 0.19 | 1.8e-05 | 2.8e-02 | wool QTL † | 13 |  |

### Selection Signatures within population groups

#### Coloration

Many selection signatures are located around genes that have been shown to be involved inhair, eye or skin color. In particular, several detected regions include candidate genes that are involved in the development and migration of melanocytes and in pigmentation : EDN3, KIT, KITLG, MC1R and MITF. For all these genes except MITF, we have quite strong evidence that they are the genes targeted by selection in the detected region. In the SWA group, EDN3 was included in the detected region for both FLK and hapFLK, and in both cases it was the closest gene to the highest test value. KIT and KITLG were both included in a detected region (with relatively few genes) for two different geographical groups, and were very close to the position with the smallest p-value in one of those. MC1R was also in a detected region for two different groups, NEU and ITA. In the two cases it was not very close to the maximum of the signal, but we note that the black skin or coat color is an important characteristic of the two populations that have been found under selection in this region, the Irish Suffolk and Sardinian Ancestral Black. This observation, together with the fact that MC1R mutations are responsible for coat color patterns in mammals (*e.g* in cattle [27]), supports the hypothesis that MC1R is a good candidate for the signatures we observed.

Although not listed in Table 2, SOX10 and ASIP, two other genes implied in pigmentation, also show some evidence of selection. In the ITA group, the q-value of hapFLK near SOX10 is 6.2% and almost reaches the significance threshold of 5%. Similarly, the two closest SNP to ASIP (*s66432* and *s12884*) present suggestive FLK p-values of respectively 7.5 10^−4^ and 6.8 10^−5^ in the ASI group, and one (*s12884*) is significantly differentiated between the ancestral groups. All these genes have previously been reported as being likely selection targets and/or associated to color patterns in different mammalian species. Finally, we found a signal for selection centered on the BNC2 gene, that has recently been associated with skin pigmentation in humans [28]. All population groups present at least one selection signature which is very likely related to one of the above genes, reflecting the widespread importance of color patterns to define sheep breeds.

Inferring a precise history of underlying causal mutations for color patterns in this dataset is hard for several reasons: the precise phenotypic characterizations of coat color patterns in the Sheep HapMap breeds are not available; the 50 K SNP array used does not offer sufficient density to associate a given selection signature to a specific set of polymorphisms; finally, from the literature, it appears that coat color is a complex trait, with high genetic heterogeneity. In particular, mutations in different genes can give rise to the same phenotype (*e.g.* in horses [29]). Also, within a gene different mutations can give rise to different phenotypes, *e.g* mutations in the MC1R gene (also named the extension locus) have been associated to a large panel of skin or coat colors [27,30,31]. Deciphering selection signatures related to coat color in sheep and in particular identifying the causal variants under selection will require sequencing these genes for individuals from several breeds with diverging color patterns. This in turn will help to understand the evolutionary history of the breeds and the effect of selection [32]. To potentially help in this task, in Table S1 we list, for each “color gene”, the populations that have likely been selected for.

#### Morphology

Another group of genes that are found within selection signatures have known effects on body morphology and development. NPR2, HMGA2 and BMP2, pointed out previously [4] are confirmed as good positional candidates by our study. We also found strong evidence for selection on WNT5A, ALX4 or EXT2, and two HOX gene clusters (HOXA and HOXC). WNT5A and ALX4 are two genes involved in the development of the limbs and skeleton. Mutations in WNT5A are causing the dominant Human Robinow syndrome, characterized by short stature, limb shortening, genital hypoplasia and craniofacial abnormalities [33]. ALX4 loss of function mutations cause polydactily in the mouse, through disregulation of the sonic hedgehog (SHH) signaling factor [34,35]. Moreover, the ALX4 protein has been shown to bind proteins from the HOXA (HOXA11 and HOXA3) and HOXC (HOXC4 and HOXC5) clusters [36]. Located just besides ALX4 and corresponding to the same selection signature, EXT2 is responsible for the development of exostose in the mouse [37]. *HOX* genes are responsible for antero-posterior development and skeletal morphology along the anterior-posterior axis in vertebrates. The selection signature around HOXA is a recent selection signature in the SWA group, while that around HOXC is an ancestral signature with a high differentiation of the ASI ancestor compared to AFR and SWA (Table 3).

Finally, we note that an ancestral selection signature is found near the ACAN gene, whose expression was shown to be upregulated by BMP2 [38], another candidate gene for selection. Three genes within the selection signature are found closer to the maximum test value than ACAN, but these are in silico predicted genes, whose protein coding function has not been confirmed, so ACAN seems to be overall a better candidate for explaining selection in the region. Mutations in the ACAN gene have been shown to induce osteochondrosis [39] and skeletal dysplasia [40]. The ACAN region has also been shown to be associated with height in humans [41].

#### Traits of agronomic importance

Sheeps have been raised for meat, milk and wool production. Under selection signatures, we found several genes associated with these production traits. In addition to the selection signature in Texels on the MSTN gene for increased muscularity [42], discussed in [13], we detected a selection signature centered on HDAC9 and including few other genes, which could also be linked to muscling. HDAC9 is a known transcriptional repressor of myogenesis. Its expression has been shown to be affected by the callypige mutation in the sheep at the DLK1-DIO3 locus [43]. The signature around HDAC9 corresponds to a selection signature in the Garut breed from Indonesia, a breed used in ram fights. As already discussed, one selection signature contains ABCG2, a gene underlying a QTL with large effects on milk production (yield and composition) in cattle [20]. Also, one of the ancestral selection signatures reaches its maximum value close to the INSIG2 gene, recently shown to be associated with milk fatty acid composition in Holstein cattle [44]. Two selection signatures could be related to wool characteristics, one in the CEU group including the FGF5 gene, partly responsible for hair type in the domestic dog [45,46], and an ancestral selection signature on chromosome 25 in a QTL region associated to wool quality traits in the sheep [47,48].

One of the strong outlying regions in the selection scan contains the PITX3 gene. Further analysis revealed that this signature was due to the German Texel population haplotype diversity differing from the other Texel samples (results not shown). It turns out that the German Texel sample consisted of a case/control study for microphtalmia [7], although the case/control status information in this sample is not given in the Sheep HapMap dataset. The consequence of such a recruitment is to bias haplotype frequencies in the region associated with the disease, which provokes a very strong differentiation signal between the German Texel and the other Texel populations. Although not related to artificial or natural selection in sheep, this signature illustrates that our method for detecting selection has the potential to identify causal variants in case/control studies, while using haplotype information.

### Ancestral signatures of selection

It is difficult to estimate how far back in time signatures of selection found in the ancestral tree took place. In particular, it would be interesting to place the divergences shown by the ancestral population tree with respect to sheep domestication. Two interesting candidate genes for ancestral selection signatures might indicate that the selection signatures captured could be rather old. First, we found selection near the TRPM8 gene, which has been shown to be a major determinant of cold perception in the mouse [49]. The pattern of allele frequency at the significant SNP (Table 3) is consistent with the climate in the geographical origins of the population groups. AFR, ASI and ITA, living in warm climates, have low frequency (0.04–0.16) of the A allele, while NEU and CEU, from colder regions, have higher frequencies (0.55–0.7), the SWE group having an intermediate frequency of 0.38. Overall, this selection signature might be due to an adaptation to cold climate through selection on a TRPM8 variant. Another selection signature lies close to a potential chicken domestication gene, TSHR [50], whose signaling regulates photoperiodic control of reproduction [51]. This selection signature was identified before [4] and our analysis indicates that selection happened before the divergence of breeds within geographic groups, consistent with an early selection event. Given its role, we can speculate that selection on the TSHR gene is related to seasonality of reproduction. Under temperate climates, sheep experience a reproductive cycle under photoperiodic control. Furthermore, there is evidence that this control was altered during domestication [52] so our analysis suggests genetic mutations in TSHR may have contributed to this alteration.

As discussed above, some of the genes found underlying ancestral selection signatures can be related to production or morphological traits (*e.g.* ASIP, INSIG2, ACAN, wool QTL), indicating that these traits have likely been important at the beginning of sheep history. The other genes that we could identify as likely selection targets in the ancestral population tree relate to immune response (GATA3) and in particular to antiviral response (TMEM154 [53], TRAF3 [54]). The most significant ancestral selection signature is centered around the NF1 gene, encoding neurofibromin. This gene is a negative regulator of the ras signal transduction pathway, therefore involved in cell proliferation and cancer, in particular neurofibromatosis. Due to this central role in intra-cellular signaling, mutations affecting this gene can have many phenotypic consequences so that its potential role in the adaptation of sheep breeds remains unclear.

## Conclusions

The Sheep HapMap dataset is an exceptional resource for sheep genetics studies. In a population genomics context, our study shows that the rich information contained in these data permits to start unraveling the genetic history of sheep populations worldwide. In order to fully exploit this information, we used recent statistical approaches that account for the relationship between populations and the linkage disequilibrium patterns (haplotype diversity). This allowed detecting with confidence more selection signatures and identifying for most of them the selected populations. Among these new selection signatures detected by our study, several result from recent selection and include good positional candidate genes with functions related to pigmentation (KITLG, EDN3), morphology (WNT5A, ALX4, EXT2, HOXA cluster) or production traits (HDAC9). Two ancestral selection signatures are also of particular interest as they harbor genes (TRPM8 and TSHR) whose functions (cold and photoperiodic perception respectively) seem highly relevant to the selection response during the early history of domestic sheep.

With information on adaptive genome regions and selected populations, we hope that our work will foster new studies to unravel the underlying biological mechanisms involved. To this aim, it is likely that further phenotypic and genetic data are required. On the genetics side, even though the SNP array used in this study was sufficient to localize genome regions harboring adaptive mutations, its density and the SNP ascertainment bias resulting from its design did not allow to tag the causative mutation precisely. Elucidating the causal variation underlying selection signatures will thus most likely require large scale sequencing data.

Genome scans for selection, including this one, are identifying regions that are outliers from a statistical model and do not require to specify an alternative hypothesis based on phenotypic records. While this can be seen as an advantage for the initial localization of genome regions, it is a limitation for the identification of biological processes involved. Gathering phenotypic records in specific populations, in particular for color and morphology traits, will be needed to go further.

## Methods

### Selecting populations and animals

Seventy-four breeds are represented in the Sheep HapMap data set, but we only used a subset of these breeds in our genome scan. We removed the breeds with small sample size (< 20 animals), for which haplotype diversity cannot be determined with sufficient precision. Based on historical information, we also removed all breeds resulting from a recent admixture or having experienced a severe recent bottleneck. Focusing on the remaining breeds, we then studied the genetic structure within each population group, in order to detect further admixture events. We performed a standardized PCA of individual based genotype data and applied the admixture software [16].

In two population groups (AFR and NEU) the different breeds were clearly separated into distinct clusters of the PCA and showed no evidence of recent admixture (Figures S1 and S2). These samples were left unchanged for the genome scan for selection. A similar pattern was observed in three other groups (ITA, SWA, ASI), except for a few outlier animals that had to be re-attributed to a different breed or simply removed (Figures S3, S4 and S5). In the two last groups (CEU and SWE), several admixed breeds were found and were consequently removed from the genome scan analysis (Figures S6 and S7).

We performed a genome scan within each group of populations listed in Table 1, with a single SNP statistic FLK [10] and its haplotype version hapFLK [13].

### Population trees

Both statistics require estimating the population tree, with a procedure described in details in [10]. Briefly, we built a population tree for each group by first calculating Reynolds’ distances between each population pair, and then applying the Neighbor Joining algorithm on the distance matrix. For each group, we rooted the tree using the Soay sheep as an outgroup. This breed has been isolated on an Island for many generations and exhibits a very strong differentiation with all the breeds of the Sheep HapMap dataset, making it well suited to be used as an outgroup.

### FLK and hapFLK genome scans

The FLK statistic was computed for each SNP within each group. The evolutionary model underlying the FLK statistic assumes that SNP were already polymorphic in the ancestral population. To consider only loci that most likely match this hypothesis, we restricted our analysis within each group to SNP for which estimated ancestral minor allele frequency *p*_0_ was above 5%. Under neutrality, the FLK statistic should follow a *χ*^2^ distribution with *n* − 1 degrees of freedom (DF), where *n* is the number of populations in the group. Overall, the fit of the theoretical distribution to the observed distribution was very good (supporting information Text S1) with the mean of the observed distribution (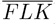) being very close to *n* − 1 (Table S3). Using 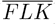 as DF for the *χ*^2^ distribution provided a better fit to the observed data than the *n* − 1 theoretical value. We thus computed FLK p-values using the *χ*^2^(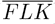) distribution. To compute the hapFLK statistic, we used of the Scheet and Stephens LD model [55], a mixture model for haplotypes which requires specifying a number of haplotype clusters to be used. To choose this number, for each group, we used the fastPHASE cross-validation based estimation of the optimal number of clusters. The results of this estimation are given in Table S2. The LD model was estimated on unphased genotype data. The hapFLK statistic is computed as an average over 20 runs of the EM algorithm to fit the LD model. As in [13], we found that the hapFLK distribution could be modeled relatively well with a normal distribution (corresponding to non outlying regions) and a few outliers; we used robust estimation of the mean and standard deviation of the hapFLK statistic to eliminate the influence of outlying (*i.e.* potentially selected) regions. This procedure was done within each group, the resulting mean and standard deviation values obtained are given in Table S2. Finally, we computed at each SNP a p-value for the null hypothesis from the normal distribution.

### Selection in ancestral groups

The within-group FLK analysis provides for each SNP an estimation of the allele frequency *p*_0_ in the population ancestral to all populations of the group. We used this information to test SNP for selection using between group differentiation, with some adjustments. First, the FLK model assumes tested polymorphisms are present in the ancestral population. SNP for which the alternate allele has been seen in only one population group are likely to have appeared after divergence (within the ancestral tree) and were therefore removed from the analysis. Second, regions selected within groups affect allele frequency in some breeds and therefore bias our estimation of the ancestral allele frequency in this group. We therefore removed all SNP that were included in within-group selection signatures. Finally, the FLK test requires a rooted population tree. For the within group analysis, we could use a very distant population to the current breeds (the Soay sheep). For the ancestral tree, we created an outgroup homozygous for ancestral alleles at all SNP.

### Identifying selected regions and candidate genes

We defined significant regions for each statistic and within each group of populations. Using the neutral distribution (*χ*^2^ for FLK and Normal for hapFLK), we computed the p-value of each statistic at each SNP. To identify selected regions, we estimated their q-value [56] to control the FDR. For FLK, SNP with a q-value below 0.1 were considered significant, which by definition implies that we expect 10% of false positives among our detected SNP. Since the power of hapFLK is greater than that of FLK [13], we used a q-value threshold of 0.05, therefore controlling FDR at the 5% level. For the FLK analysis in ancestral populations, we used an FDR threshold of 5%.

We then aimed at identifying genes that seem good candidates for explaining selection signatures. We proceeded differently for the single SNP FLK and hapFLK. For FLK, we considered that significant SNP less than 500Kb apart were capturing the same selection signal. Then, we considered as potential candidate genes any gene that lies less than 1Mb of any significant SNP. For hapFLK, the genome signal is much more continuous than single SNP tests, because the statistic captures multipoint LD with the selected mutations. A consequence is that the significant regions can span large chromosome intervals. To restrict the list of potential candidate genes, and target only the ones closest to the most significant SNP, we restricted our search to the part of the signal where the difference in hapFLK value with the most significant SNP was less than 0.5*σ*. This allowed taking into consideration the profile of the hapFLK signal, *i.e.* if the profile resembles a plateau, the candidate region will be rather broad while very sharp hapFLK peaks will provide a narrower candidate region. We extracted all protein coding genes present in the significant regions using the Ensembl Biomart tool (http://www.ensembl.org/biomart/) for Ovis Aries 3.1 genome assembly. These full lists are provided as Supplementary data (Supporting Dataset 1 and Supporting Dataset 2). Within each candidate region, genes were ranked according to their distance from the most significant position of the region (the larger the rank, the larger the distance). The functional candidate genes shown in Table 2 and discussed in the manuscript were chosen based on this rank and / or on their implication in previous association or sweep detection studies.

## Acknowledgments

Data analyses were performed on the computer cluster of the bioinformatics platform Toulouse Midi-Pyrenees. We would like to thank Wendy Brand-Williams for her careful reading of the manuscript.

The members of the International Sheep Genomics Consortium who contributed samples and expertise towards the design and execution of ovine SNP50k genotyping and next generation sequencing, and/or coauthors of [4] include: Juan Jose Arranz, Universidad de Leon; Georgios Banos, Aristotle University of Thessaloniki; William Barendse, CSIRO Livestock Industries; Ahmedn El Beltagy, Animal Production Research Institute; Jorn Benenwitz, University of Hohenheim; Steven Bishop, The Roslin Institute; Simon Boitard, INRA; Lutz Bunger, Scottish Agricultural College; Jorge Calvo, CITA; Antonello Carta, Agris Sardegna; Ibrahim Cemal, Adnan Menderes University; Elena Ciani, University of Bari; Noelle Cockett, University of Utah; Dave Coltman, University of Alberta; Brian Dalrymple, CSIRO Livestock Industries; Mariasilvia DAndrea, Universit degli Studi del Molise; Ottmar Distl, University of Veterinary Medicine Hannover; Cord Drogemuller, Institute of Genetics, University of Berne; Georg Erhardt, Institut fr Tierzucht und Haustiergenetik Justus-Liebig-Universitt Gieen; Emma Eythorsdottir, Agricultural University of Iceland; Kimberly Gietzen, Illumina Inc.; Clare Gill, Texas A& M University; Elisha Gootwine, The Volcani Center; Vidya Gupta, National Chemical Laboratory; Olivier Hanotte, University of Nottingham; Ben Hayes, Department of Primary Industries Victoria; Michael Heaton, USDA MARC; Stefan Hiendleder, University of Adelaide; Han Jialin, ILRI and CAAS; Juha Kantanen, MTT Agrifood Research; Matthew Kent, CiGene; James Kijas, CSIRO Livestock Industries; Denis Larkin, University of Aberystwyth; Johannes A. Lenstra, Utrecht University; Kui Li, Lhasa People Hospital, Tibet; Terry Longhurst, Meat and Livestock Australia; Runlin Ma, Chinese Academy of Science; Russell McCulloch, CSIRO Livestock Industries; David MacHugh, University College Dublin; Sean McWilliam, CSIRO Livestock Industries; John McEwan, AgResearch; Jillian Maddox, University of Melbourne; Massoud Malek, IAEA; Faruque Mdomar, Bangladesh Agriculture University; Despoina Miltiadou, Cyprus University of Technology; Luis V. Monteagudo Ibez, Universidad de Zaragoza; Carole Moreno, INRA; Frank Nicholas, University of Sydney; Kristen Nowak, University of Western Australia; V. Hutton Oddy, University of New England; Samuel Paiva, Embrapa; Varsha Pardeshi, National Chemical Laboratory; Josephine Pemberton, University of Edinburgh; Fabio Pilla, Universit degli Studi del Molise; Laercio R. Porto Neto, CSIRO Livestock Industries; Herman Raadsma, University of Sydney; Cyril Roberts, Caribbean Agricultural Research and Development Institute; Magali San Cristobal, INRA; Tiziana Sechi, Agris Sardegna; Paul Scheet, University of Texas M. D. Anderson Cancer Center; Bertrand Servin, INRA; Mohammad Shariflou, University of Sydney; Pradeepa Silva, University of Peradeniya; Henner Simianer, University of Goettingen; Jon Slate, University of Sheffield; Mikka Tapio, MTT; and Selina Vattathil, University of Texas M. D. Anderson Cancer Center; Vicki Whan, CSIRO Livestock Industries.

## References

1. Andersson L (2012) How selective sweeps in domestic animals provide new insight into biological mechanisms. J Intern Med 271: 1–14.

2. Zeder MA (2008) Domestication and early agriculture in the mediterranean basin: Origins, diffusion, and impact. Proceedings of the National Academy of Sciences 105: 11597–11604.

3. Kijas JW, Townley D, Dalrymple BP, Heaton MP, Maddox JF, et al. (2009) A genome wide survey of SNP variation reveals the genetic structure of sheep breeds. PLoS One 4: e4668.

4. Kijas JW, Lenstra JA, Hayes B, Boitard S, Porto Neto LR, et al. (2012) Genome-wide analysis of the world's sheep breeds reveals high levels of historic mixture and strong recent selection. PLoS Biol 10: e1001258.

5. Garcia-Gámez E, Reverter A, Whan V, McWilliam SM, Arranz JJA, et al. (2011) Using regulatory and epistatic networks to extend the findings of a genome scan: identifying the gene drivers of pigmentation in merino sheep. PLoS ONE 6: e21158.

6. Moradi MH, Nejati-Javaremi A, Moradi-Shahrbabak M, Dodds KG, McEwan JC (2012) Genomic scan of selective sweeps in thin and fat tail sheep breeds for identifying of candidate regions associated with fat deposition. BMC Genet 13: 10.

7. Becker D, Tetens J, Brunner A, Burstel D, Ganter M, et al. (2010) Microphthalmia in Texel sheep is associated with a missense mutation in the paired-like homeodomain 3 (PITX3) gene. PLoS One 5: e8689.

8. Zhang L, Mousel MR, Wu X, Michal JJ, Zhou X, et al. (2013) Genome-Wide Genetic Diversity and Differentially Selected Regions among Suffolk, Rambouillet, Columbia, Polypay, and Targhee Sheep. PLoS One 8: e65942.

9. Excoffier L, Hofer T, Foll M (2009) Detecting loci under selection in a hierarchically structured population. Heredity (Edinb) 103: 285–98.

10. Bonhomme M, Chevalet C, Servin B, Boitard S, Abdallah J, et al. (2010) Detecting selection in population trees: the Lewontin and Krakauer test extended. Genetics 186: 241–62.

11. Bierne N, Roze D, Welch JJ (2013) Pervasive selection or is it? why are fst outliers sometimes so frequent? Molecular Ecology 22: 2061–2064.

12. Gnther T, Coop G (2013) Robust identification of local adaptation from allele frequencies. Genetics 195: 205–220.

13. Fariello MI, Boitard S, Naya H, SanCristobal M, Servin B (2013) Detecting signatures of selection through haplotype differentiation among hierarchically structured populations. Genetics 193: 929–41.

14. Lewontin RC, Krakauer J (1973) Distribution of gene frequency as a test of the theory of theselective neutrality of polymorphisms. Genetics 74: 175–195.

15. Pritchard JK, Stephens M, Donnelly P (2000) Inference of population structure using multilocus genotype data. Genetics 155: 945–59.

16. Alexander DH, Novembre J, Lange K (2009) Fast model-based estimation of ancestry in unrelated individuals. Genome Res 19: 1655–64.

17. Johnston SE, McEwan JC, Pickering NK, Kijas JW, Beraldi D, et al. (2011) Genome-wide association mapping identifies the genetic basis of discrete and quantitative variation in sexual weaponry in a wild sheep population. Mol Ecol 20: 2555–66.

18. Dominik S, Henshall JM, Hayes BJ (2012) A single nucleotide polymorphism on chromosome 10 ishighly predictive for the polled phenotype in Australian Merino sheep. Anim Genet 43: 468–70.

19. Akey JM, Ruhe AL, Akey DT, Wong AK, Connelly CF, et al. (2010) Tracking footprints of artificial selection in the dog genome. Proc Natl Acad Sci U S A 107: 1160–5.

20. Cohen-Zinder M, Seroussi E, Larkin DM, Loor JJ, Everts-van der Wind A, et al. (2005) Identification of a missense mutation in the bovine ABCG2 gene with a major effect on the QTL on chromosome 6 affecting milk yield and composition in Holstein cattle. Genome Res 15: 936–44.

21. Eberlein A, Takasuga A, Setoguchi K, Pfuhl R, Flisikowski K, et al. (2009) Dissection of genetic factors modulating fetal growth in cattle indicates a substantial role of the non-smc condensin i complex, subunit g (ncapg) gene. Genetics 183: 951–964.

22. Bongiorni S, Mancini G, Chillemi G, Pariset L, Valentini A (2012) Identification of a short region on chromosome 6 affecting direct calving ease in piedmontese cattle breed. PLoS ONE 7: e50137.

23. Mancini G, Gargani M, Chillemi G, Nicolazzi EL, Marsan PA, et al. (2014) Signatures of selection in five Italian cattle breeds detected by a 54 K SNP panel. Mol Biol Rep 41: 957–65.

24. Rothammer S, Seichter D, Forster M, Medugorac I (2013) A genome-wide scan for signatures of differential artificial selection in ten cattle breeds. BMC Genomics 14: 908.

25. Pintus E, Sorbolini S, Albera A, Gaspa G, Dimauro C, et al. (2013) Use of locally weighted scatterplot smoothing (LOWESS) regression to study selection signatures in Piedmontese and Italian Brown cattle breeds. Anim Genet.

26. Druet T, Perez-Pardal L, Charlier C, Gautier M (2013) Identification of large selective sweeps associated with major genes in cattle. Anim Genet 44: 758–62.

27. Klungland H, Våge DI, Gomez-Raya L, Adalsteinsson S, Lien S (1995) The role of melanocyte-stimulating hormone (MSH) receptor in bovine coat color determination. Mammalian genome 6: 636–9.

28. Jacobs LC, Wollstein A, Lao O, Hofman A, Klaver CC, et al. (2013) Comprehensive candidate gene study highlights UGT1A and BNC2 as new genes determining continuous skin color variation in Europeans. Hum Genet 132: 147–58.

29. Hauswirth R, Haase B, Blatter M, Brooks Sa, Burger D, et al. (2012) Mutations in MITF and PAX3 cause “splashed white” and other white spotting phenotypes in horses. PLoS genetics 8:e1002653.

30. Lin JY, Fisher DE (2007) Melanocyte biology and skin pigmentation. Nature 445: 843–50.

31. Joerg H, Fries HR, Meijerink E, Stranzinger GF (1996) Red coat color in Holstein cattle is associated with a deletion in the MSHR gene. Mammalian genome 7: 317–8.

32. Linnen CR, Poh YP, Peterson BK, Barrett RDH, Larson JG, et al. (2013) Adaptive evolution of multiple traits through multiple mutations at a single gene. Science 339: 1312–6.

33. Person AD, Beiraghi S, Sieben CM, Hermanson S, Neumann AN, et al. (2010) WNT5A mutations in patients with autosomal dominant Robinow syndrome. Developmental dynamics 239: 327–37.

34. Kuijper S, Feitsma H, Sheth R, Korving J, Reijnen M, et al. (2005) Function and regulation of Alx4 in limb development: complex genetic interactions with Gli3 and Shh. Developmental biology 285: 533–44.

35. Qu S, Tucker SC, Ehrlich JS, Levorse JM, Flaherty LA, et al. (1998) Mutations in mouse Aristaless-like4 cause Strong's luxoid polydactyly. Development 125: 2711–21.

36. Ravasi T, Suzuki H, Cannistraci CV, Katayama S, Bajic VB, et al. (2010) An atlas of combinatorial transcriptional regulation in mouse and man. Cell 140: 744–752.

37. Stickens D, Zak BM, Rougier N, Esko JD, Werb Z (2005) Mice deficient in Ext2 lack heparan sulfate and develop exostoses. Development 132: 5055–68.

38. Noguchi Ki, Watanabe Y, Fuse T, Takizawa M (2010) A new chondrogenic differentiation initiator with the ability to up-regulate sox trio expression. Journal of Pharmacological Sciences 112: 89–97.

39. Stattin EL, Wiklund F, Lindblom K, Onnerfjord P, Jonsson BA, et al. (2010) A missense mutation in the aggrecan C-type lectin domain disrupts extracellular matrix interactions and causes dominant familial osteochondritis dissecans. American journal of human genetics 86: 126–37.

40. Tompson SW, Merriman B, Funari Va, Fresquet M, Lachman RS, et al. (2009) A recessive skeletal dysplasia, SEMD aggrecan type, results from a missense mutation affecting the C-type lectin domain of aggrecan. American journal of human genetics 84: 72–9.

41. Weedon MN, Lango H, Lindgren CM, Wallace C, Evans DM, et al. (2008) Genome-wide association analysis identifies 20 loci that influence adult height. Nature genetics 40: 575–83.

42. Clop A, Marcq F, Takeda H, Pirottin D, Tordoir X, et al. (2006) A mutation creating a potential illegitimate microRNA target site in the myostatin gene affects muscularity in sheep. Nat Genet 38: 813–8.

43. Fleming-Waddell JN, Olbricht GR, Taxis TM, White JD, Vuocolo T, et al. (2009) Effect of DLK1 and RTL1 but not MEG3 or MEG8 on muscle gene expression in Callipyge lambs. PloS one 4:e7399.

44. Rincon G, Islas-Trejo A, Castillo AR, Bauman DE, German BJ, et al. (2012) Polymorphisms in genes in the SREBP1 signalling pathway and SCD are associated with milk fatty acid composition in Holstein cattle. J Dairy Res 79: 66–75.

45. Housley DJE, Venta PJ (2006) The long and the short of it: evidence that fgf5 is a major determinant of canine hair-itability. Animal Genetics 37: 309–315.

46. Cadieu E, Neff MW, Quignon P, Walsh K, Chase K, et al. (2009) Coat variation in the domestic dog is governed by variants in three genes. Science 326: 150–3.

47. Ponz R, Moreno C, Allain D, Elsen JM, Lantier F, et al. (2001) Assessment of genetic variation explained by markers for wool traits in sheep via a segment mapping approach. Mamm Genome 12: 569–72.

48. Bidinost F, Roldan D, Dodero A, Cano E, Taddeo H, et al. (2007) Wool quantitative trait loci in merino sheep. Small Ruminant Research 74: 113–118.

49. Bautista DM, Siemens J, Glazer JM, Tsuruda PR, Basbaum AI, et al. (2007) The menthol receptor TRPM8 is the principal detector of environmental cold. Nature 448: 204–8.

50. Rubin CJ, Zody MC, Eriksson J, Meadows JRS, Sherwood E, et al. (2010) Whole-genome resequencing reveals loci under selection during chicken domestication. Nature 464: 587–91.

51. Nakao N, Ono H, Yamamura T, Anraku T, Takagi T, et al. (2008) Thyrotrophin in the pars tuberalis triggers photoperiodic response. Nature 452: 317–22.

52. Balasse M, Tresset A (2007) Environmental constraints on the reproductive activity of domestic sheep and cattle : what latitude for the herder? Anthropozoologica 42: 71–88.

53. Heaton MP, Clawson ML, Chitko-Mckown CG, Leymaster Ka, Smith TPL, et al. (2012) Reduced lentivirus susceptibility in sheep with TMEM154 mutations. PLoS genetics 8: e1002467.

54. Oganesyan G, Saha SK, Guo B, He JQ, Shahangian A, et al. (2006) Critical role of TRAF3 in the Toll-like receptor-dependent and -independent antiviral response. Nature 439: 208–11.

55. Scheet P, Stephens M (2006) A fast and flexible statistical model for large-scale population genotype data: applications to inferring missing genotypes and haplotypic phase. Am J Hum Genet 78: 629–44.

56. Storey JD, Tibshirani R (2003) Statistical significance for genomewide studies. Proc Natl Acad Sci U S A 100: 9440–5.

